# Physically intelligent insect-inspired antenna sensors enhance tactile feature perception by active touch

**DOI:** 10.1101/2025.10.20.683587

**Authors:** Lingsheng Meng, Parker McDonnell, Kaushik Jayaram, Jean-Michel Mongeau

## Abstract

Soft robotic sensors today struggle to interpret complex tactile scenes without incurring significant computational costs. Inspired by insect antennae—compliant, distributed sensors that efficiently process tactile information through physical intelligence—we investigated whether mechanical design and active touch sensing strategies could enhance robotic tactile feature perception. We hypothesized that insect-inspired antenna dynamics, specifically flexural stiffness gradients and active touch speed, could simplify tactile classification. Using a sim-to-real framework that bridges bioinspired computational models with a multi-link soft robot antenna, we introduce the notion of tactile fields—spatiotemporal representations of tactile stimuli shaped by contact location, feature type, and active touch speed. Our analyses show that cockroach-inspired antenna mechanics jointly with active touch speeds improve feature classification accuracy compared to conventional sensors with uniform flexural stiffness gradient by increasing tactile data sparsity and dispersion. An exploration of stiffness and damping of antenna mechanics revealed design trade-offs that influence tactile discrimination and structural stability. Through sim-to-real transfer, stiffness gradients and structured active touch motions were demonstrated on a miniature distributed soft robotic antenna, validating their effectiveness in real-world robotic systems. Taken together, this work presents a biologically grounded framework for tactile sensor design that reduces computational load and enhances adaptability.

## Introduction

A critical challenge in robotics is in efficiently interpreting tactile information from complex environments. Animals, particularly insects, excel at navigating dynamic, cluttered habitats by leveraging physical intelligence—the intrinsic ability of their body mechanics to simplify sensory processing and reduce cognitive load [1,2]. While robotic perception through non-contact modalities such as vision has significantly advanced, tactile sensors inspired by biological systems like mammalian whiskers [3], lobster antennae [4,5], and insect antennae [6–9] remain relatively less developed, particularly at the scale of insects. Unlike vision, tactile sensing which results from physical interactions with the environment, provides unique insights into object geometry, texture, and mechanical properties. However, these physical interactions pose challenges in terms of sensor design, information processing and perception. Additionally, active sensing behaviors that structure sensor motion in synergy with sensor mechanics likely modulate tactile sensitivity and classification accuracy [10]. For example, animals might adjust their exploration speeds to optimize tactile perception, analogous to the speed-accuracy trade-offs during decision-making tasks extensively studied in vision [11–13]. Slower contact speeds potentially allow extended sensory integration times, increasing the quality and resolution of tactile information (**Figure 1A,B**). Despite these expected benefits, the relationship between tactile exploration speeds and the fidelity of feature discrimination has yet to be quantified systematically in insect tactile systems. Addressing this gap would reveal how active sensing modulates tactile perception, providing insights into fundamental trade-offs between sensory accuracy and energetic costs in insects, and informing the design of energy-efficient robotic tactile sensors. Investigating and adopting biological principles of physical intelligence could therefore lead to advancements in robotic sensing, enhancing efficiency, adaptability, and scalability.

**Figure 1.**
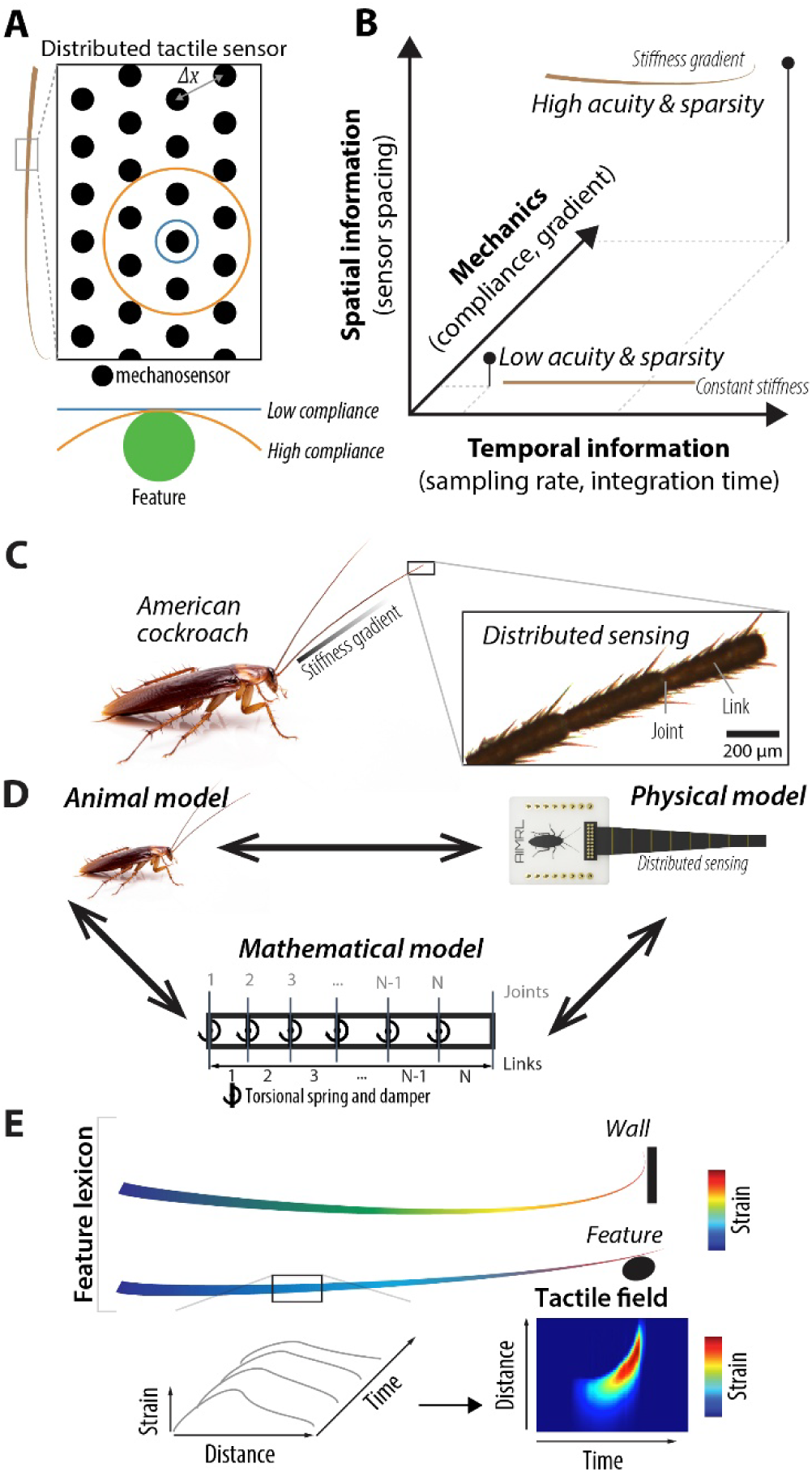
An insect-inspired design framework for tactile perception with distributed sensing. (**A**) A high compliance distributed sensor increases the information available by activating more mechanosensors. Δ*x*: sensor spacing. (**B**) Tactile sensation with distributed sensors is influenced by spatial information, temporal information and mechanics. (**C**) The American cockroach *Periplaneta americana* has a pair of compliant, distributed antennae for touch sensation. (**D**) Framework combining animal experiments, mathematical and physical modeling. (**E**) Contact of the antenna with features generates a spatiotemporal gradient of strain along its length. The relationship between strain, distance and time can be represented as a tactile field.

Insect antennae exemplify these principles through their distributed mechanosensors and adaptive mechanical properties, such as flexural stiffness gradients (henceforth referred to as stiffness gradients), enabling rapid and precise tactile perception and guidance [14,15]. Such structural features may facilitate a diverse range of tasks from rapid obstacle avoidance during locomotion to fine-scale tactile exploration [16,17]. Previous studies in mammalian whiskers and insect wings have established that nonuniform stiffness gradients can improve spatial acuity and simplify sensory processing by mechanically mediating sensory input [18–20]. Furthermore, physical antenna sensors with stiffness gradients demonstrated that tapered antennae facilitate tactile acuity by increasing the spatial resolution of tactile sensing and mitigating mechanical perturbations (**Figure 1A**) [6,15,21]. However, while the importance of stiffness gradient is well-recognized, precisely how these mechanical adaptations influence tactile feature perception—by potentially enhancing data sparsity, feature separability, and reducing cognitive demand remains poorly understood (**Figure 1B**). Specifically, we develop a systematic framework to explore how stiffness gradients influence tactile feature discrimination, such as distinguishing shapes, sizes, and contact locations.

Cockroach antennae represent ideal biological systems for testing these hypotheses. The American cockroach (*Periplaneta americana*) is a touch specialist that relies on its antennae—which is approximately 1.5 times its body length—for feature discrimination during tactile exploration acquiring critical information about feature position, texture, contact angle, and distance [17,22–24](**Figure 1C**). Their antennae have a distinct mechanical profile characterized by an exponentially decreasing stiffness gradient [15], a feature shared with other biological tactile sensors [25], potentially hinting at convergent evolution-led design principle. However, unlike mammalian whiskers, insect antennae feature a large population of mechanosensors distributed throughout their length. For example, each cockroach antenna has roughly 40,000 mechanosensors distributed along its length [26,27]. Furthermore, it remains unclear how cockroaches precisely integrate and interpret signals from thousands of mechanosensors to discriminate tactile features and guide adaptive behaviors. One possibility is that mechanical properties of the antennae, such as intrinsic stiffness gradients, in synergy with active behavioral strategies, such as preferred contact speeds, may simplify the sensory encoding process, enhance tactile discrimination and reduce cognitive demand.

In this work, we present and validate a bioinspired computational framework to study how sensor mechanics and dynamics shape tactile information via tactile fields and to inspire robotic tactile sensors. Using detailed computational simulations grounded in recent morphological and mechanical characterization of cockroach antennae [28], we systematically examined how antenna stiffness gradients and contact speed shape tactile sensory data to influence tactile feature discrimination (**Figure 1D**). We generated broad datasets of tactile fields—spatiotemporal representations of mechanical stimuli (**Figure 1E**)—and employed dimensionality reduction and classification analyses. Our results demonstrate that cockroach-inspired antenna mechanics and slower active sensing speeds substantially increase tactile data sparsity and dispersion, enhancing tactile feature classification compared to conventional mechanics, e.g., a uniform stiffness gradient. We further explore the parameter space of the cockroach-inspired antenna mechanics to answer the question: how do base stiffness, exponential decay, and damping-to-stiffness ratio collectively shape the tactile fields? To validate the generality of our framework, we performed a sim-to-real transfer using a miniature, distributed soft robotic antenna. By comparing the classification performance of this robotic antenna against its simulated counterpart, we demonstrate that our framework accurately captures the interplay between structural mechanics and active sensing strategies. Collectively, this study established a robust, scalable tool for the design of tactile sensors, providing a platform to explore how physical intelligence can be transformed into robotic tactile sensing technologies, enabling more efficient and adaptive robotic systems.

## Results

### Active touch interactions with features generate diverse tactile fields

Animals and robots routinely encounter mechanical features with finite span, such as post-like features from plant or grass stalks, obstacles such as pebbles, features from conspecifics or predators, etc. To systematically investigate how distributed mechanosensory probes like antennae mechanically represent such environmental features, we examined how distinct mechanical contacts—varying by speed, location, and object geometry—generate sensory information encoded as tactile fields. Using a computational model inspired by cockroach antenna mechanics (**Figure 2A–C**; **Movie S1**) which enabled the high-throughput generation of tactile fields (**Movie S2**), we explored how antenna mechanics might simplify distributed information processing during contact with a small-field feature or to estimate distance to a wide-field feature, i.e. a wall. First, our model confirmed that antenna mechanics can simplify distance estimation to wide-field tactile features (**Figure S1**, **Movie S3**, see Supplement). To create a controlled dataset of tactile interactions relevant to antennal sensing during locomoting, we simulated mechanical loading across a range of contact conditions. Specifically, we defined 90 distinct tactile stimuli across three contact speeds, five contact locations, and six contact features (**Figure 2D**; see Materials and Methods). The contact features included two different sizes and three shapes. The shapes were chosen to approximate a point contact (“triangle”) and distributed contacts of distinct curvatures (“box” and “ball”), which capture a range of contact scenarios that enable controlled comparisons across conditions. Contact with these features generated a broad set of tactile fields sensitive to size, shape and contact speed (**Figure 2E**). Qualitatively, the resulting tactile fields appear distinctive as they demonstrated considerable variability depending on feature type, location of contact, and speed of interaction, suggesting that mechanical properties of the antenna inherently structure sensory data. Thus, the tactile field dataset provides a comprehensive basis for quantitatively analyzing how insect-inspired antenna mechanics and behavioral modulation via contact speed may simplify tactile information processing as detailed below.

**Figure 2.**
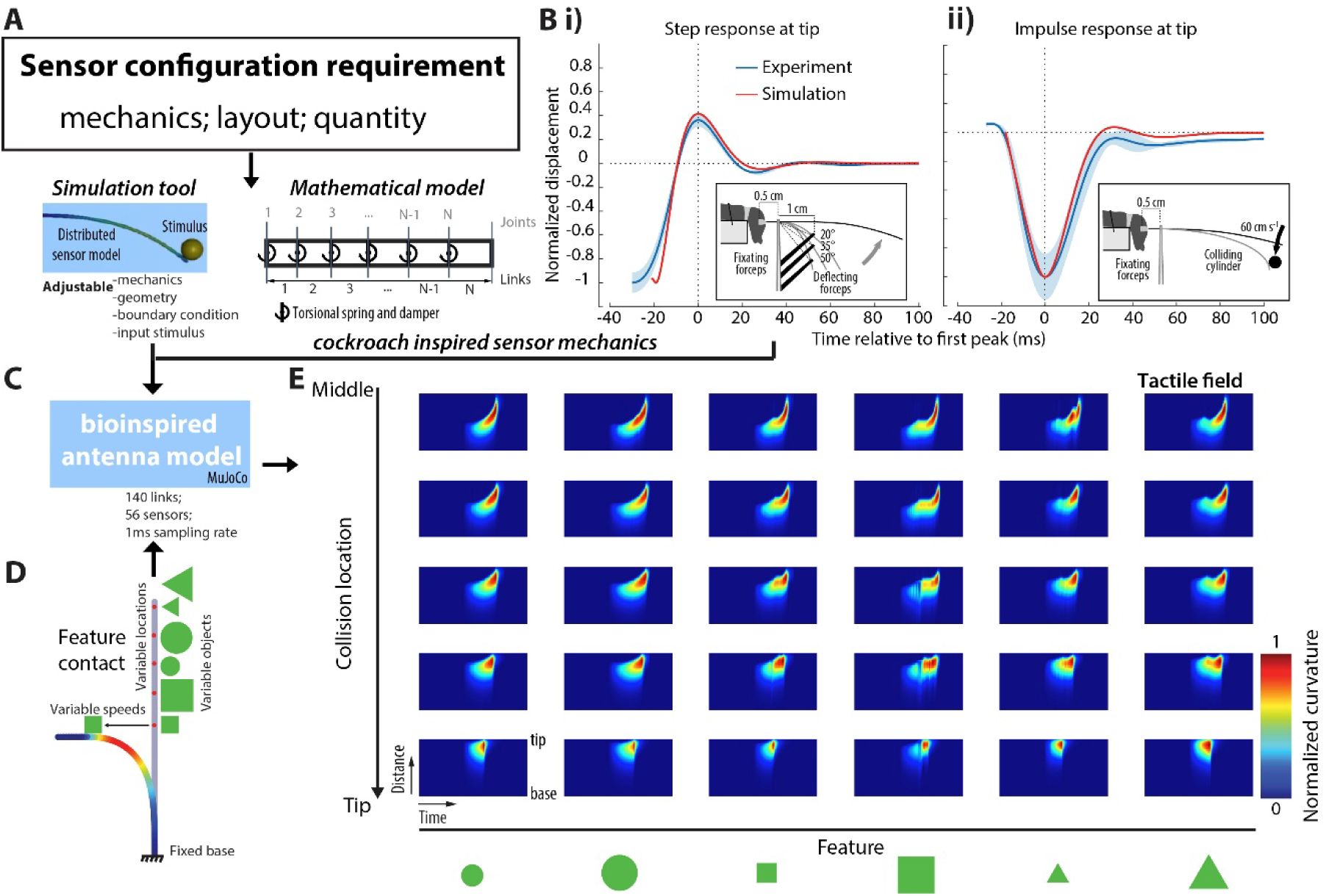
A simulation-based framework for designing distributed sensors with mechanical intelligence. (**A**) Simulation tool for distributed sensor design. A modular simulation framework computes mechanical response and sensor configuration using input stimuli and adjustable mechanical parameters. (**B**) Model validation. Tip displacement in both experiments (blue) and simulations (red) from step and impulse tests. Data from *n* = 5 animals in each scenario. Line: mean. Shaded region: ±1 STD. Adapted from [15]. (**C**) Bioinspired application. A distributed sensor array embedded in a segmented antenna model captures mechanical intelligence through biologically tuned properties. (**D**) Mechanical input variation. Six distinct features (green) impacted the antenna at five locations from middle to tip (red dots), simulating diverse tactile interactions. (**E**) Tactile field outputs. Spatiotemporal tactile fields from feature contacts under slow collision speed demonstrate sensitivity across the antenna with bioinspired sensor mechanics.

### Sparse tactile bases encode distinct feature locations

To uncover how insect-inspired antenna mechanics might influence tactile feature encoding and identify common features across all tactile fields, we formulated the following efficient encoding of tactile fields based on convolutive non-negative matrix factorization (cNMF; see Materials and Methods). This scheme decomposed the tactile fields into a set of spatiotemporal bases and their corresponding activations. To evaluate efficiency, i.e., the influence of the number of bases, *w*_*i*_(*x*, *t*), on tactile field encoding, we varied the number of bases, *M*, and obtained the bases along with their corresponding activations. Increasing the number of bases reduced the reconstruction error (L2 norm residual), indicating improved fidelity of the representation at the cost of increased dimensionality (**Figure 3B**). We found that similar patterns with clear spatial and temporal organizations emerged when the number of bases, *M*, was varied (**Figure 3A**). For instance, in the case of three bases, the first basis exhibited the highest activity in the lower left corner of the tactile field, indicating its activation during early time points and in the proximal region of the antenna. The second basis was most active around the middle region of the antenna and spanned the entire time. The third basis was most active around the upper right corner, corresponding to later time points and the distal region of the antenna. Accordingly, contact at different locations along the antenna produced distinct activation patterns across the bases: collisions near the proximal end primarily activated the first basis, while collisions at the tip strongly engaged the third basis (**Figure 3C**). While it is known that antenna bending is localized near the point of contact [15], this decomposition reveals a more compact representation: a continuum of possible contact locations can be encoded using a small number of structured bases. These sparse bases obtained through cNMF efficiently encode and preserve mechanical stimulus information; specifically, bases represent distinct contact locations, which could facilitate tactile feature discrimination. We repeated these tests for two different feature shapes (ball and box) and obtained similarly sparse bases. Collectively, these results indicate that antenna mechanics inherently structure tactile information into distinct spatiotemporal bases aligned with contact locations. This low-dimensional encoding suggests a potential mechanism by which antenna mechanics transform high-dimensional tactile inputs into a compact representation for downstream classification.

**Figure 3.**
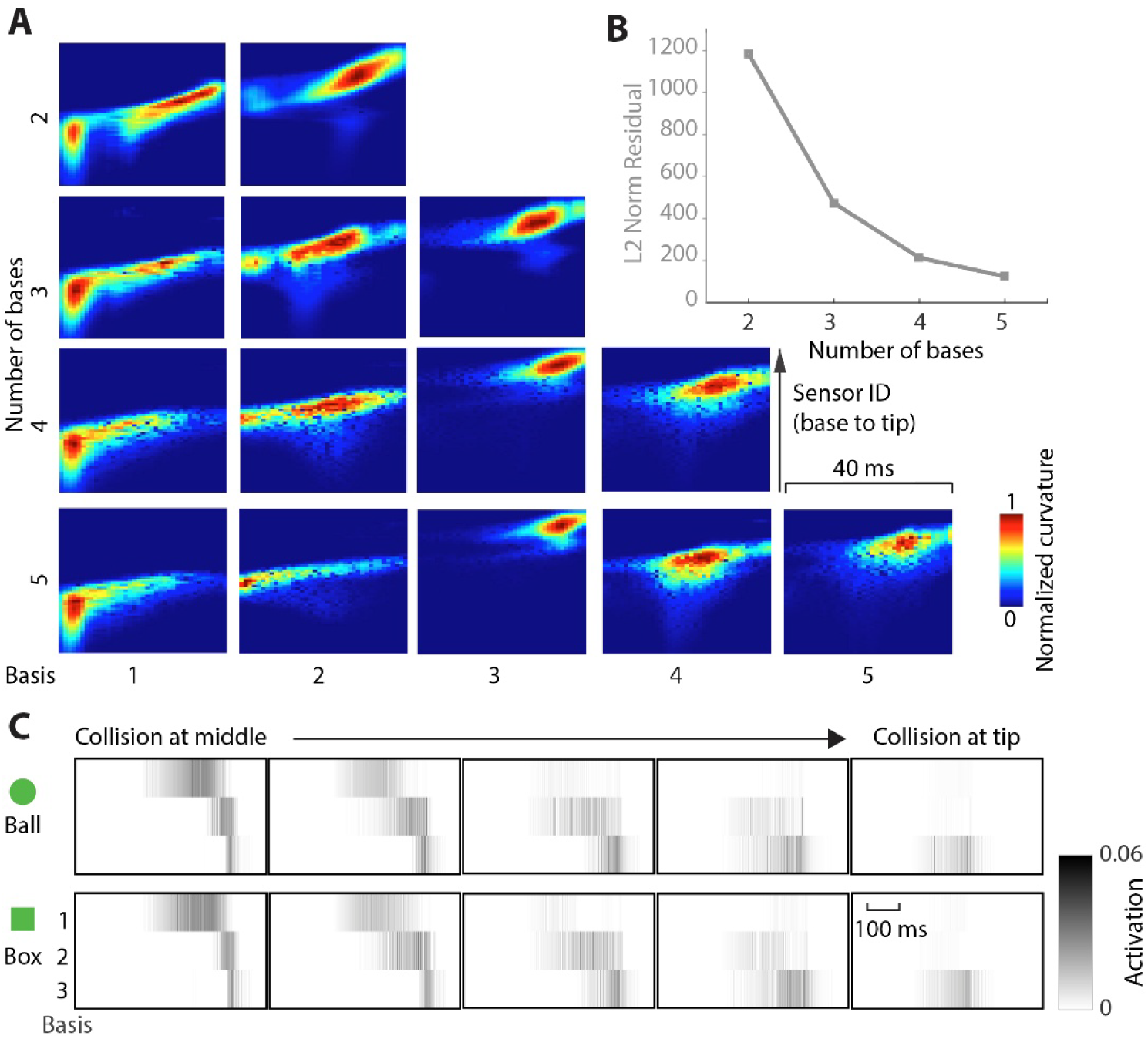
**Sparse bases represent a set of tactile features linked to contact location**. (**A**) The optimal bases from convolutive non-negative matrix factorization with different number of bases. Each row corresponds to an encoding of fixed rank from 2 to 5. The y-axis of each basis tactile field represents 56 sensors from the base to the tip of the antenna. The x-axis represents the duration of each basis (40 ms). (**B**) As the rank (*M*) increased, the encoding residual decreased. (**C**) Example activation vectors of fixed rank *M* = 3 for the ball and box collisions.

### Slower active touch speeds enhance tactile feature discrimination

Given the potential importance of active behaviors on sensing, we next explored how variations in contact speed could influence tactile feature encoding and classification performance by evaluating tactile field data dispersion (see Materials and Methods). Specifically, based on the generated tactile field set, we hypothesized that a slower contact speed (across all feature types) could facilitate tactile feature classification, analogous to increasing the exposure time or active perception on a camera or eye, respectively. For instance, for the human eye, longer exposure time to a scene allows more light capture (and temporal integration), thus facilitating visual discrimination during perceptual tasks.

To assess tactile discrimination under different contact speeds, we divided the tactile field dataset and corresponding activation dataset into three groups according to contact speed: slow (0.06 m/s), medium (0.3 m/s), and fast (0.6 m/s) (**Figure 4A**). The slow contact speed was determined by measuring antenna position during active touch exploration behavior (see Materials and Methods and Supplement), whereas the fast speed corresponds to rapid running speeds during wall following [29]. We then quantified dataset dispersion within each group using multiple indices: Pearson correlation, structural similarity index measure (SSIM), mean squared error (MSE), and normalized mutual information (NMI). A higher dispersion index indicates a more distinct dataset, which should facilitate classification by increasing between-class variance. Lower contact speeds produced more dispersed datasets, as Pearson correlation, SSIM, and NMI increased with increasing contact speed in both tactile field and activation datasets, while MSE decreased (**Figure 4A** and **Figure S2**). A Kruskal-Wallis test on NMI indicated a significant effect of contact speed on both activation dataset (*p* = 2.2e-9) and tactile field dataset (*p* = 2.4e-10). Post-hoc multiple comparisons showed that all pairwise differences between speeds on both activation dataset and tactile field dataset were significant (*p*<0.05). Lower Pearson correlation, SSIM, and NMI, along with higher MSE, indicate greater variability in the dataset. Thus, slower contact speeds produced more dispersed datasets, which in turn could enhance classification performance.

**Figure 4.**
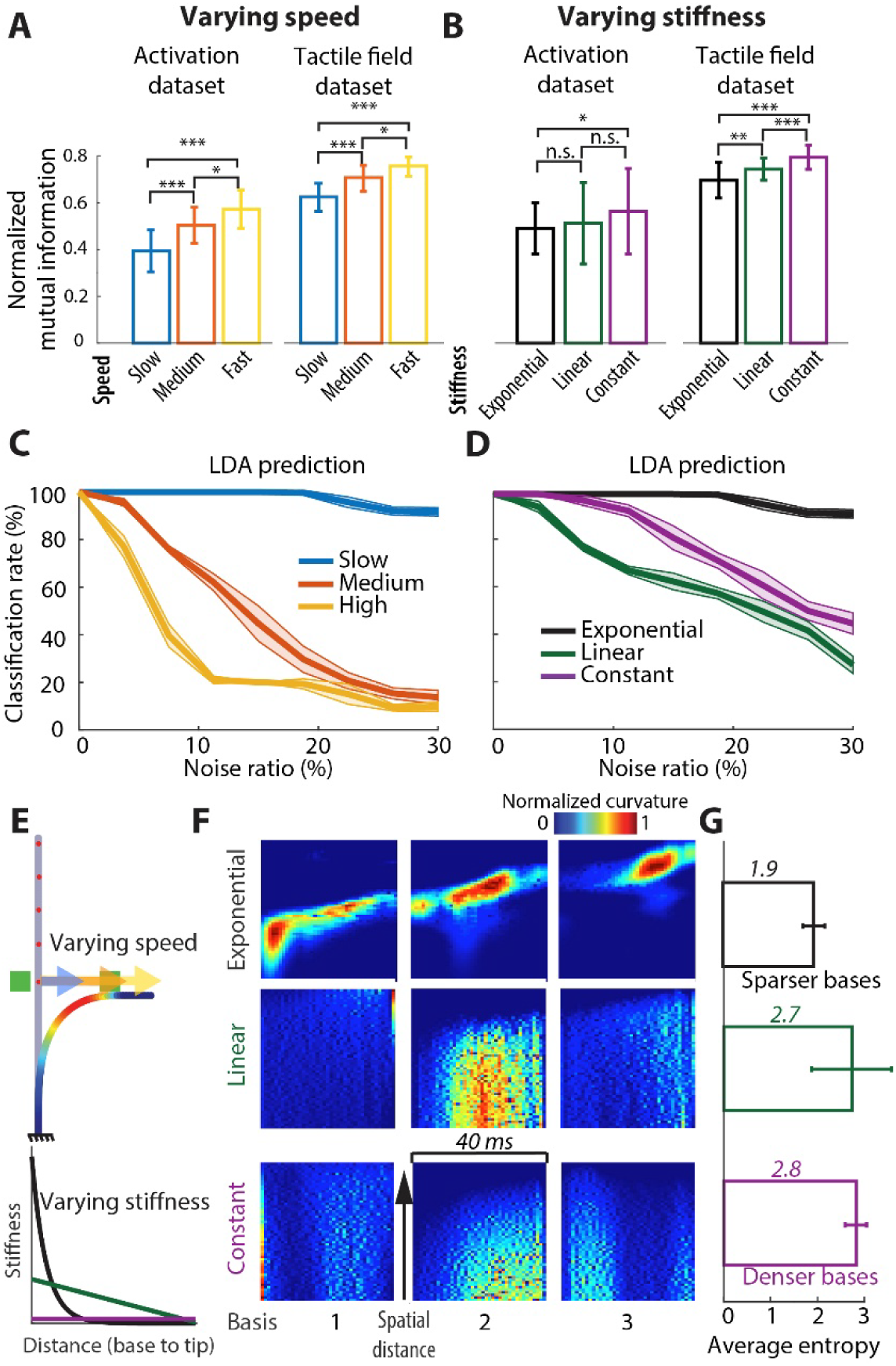
Slower contact speed and an insect-inspired antenna stiffness gradient enhance tactile feature classification. (**A**) Dispersion index (NMI, see Supplement for the other 3 indices) of three subgroups based on contact speed, shown separately for the activation dataset (left) and the tactile field dataset (right). (**B**) Dispersion index for the activation dataset (left) and the tactile field dataset (right) across three stiffness profile models: exponential, linear, and constant. (**C**) Classification rates under different levels of noise for slow, medium, and high-speed contacts. Line: mean. Shaded region: ±1 STD. (**D**) Classification rates under different levels of noise for slow-speed contacts with exponential decreasing, linear decreasing, and constant stiffness profiles. (**E**) Simulated conditions with distinct speeds and stiffness. (**F**) The optimal bases from cNMFsc with a rank of 3. (**G**) The average entropy of these bases across exponential, linear and constant flexural stiffness. Error bars: mean ± 1 standard deviation.

To test this prediction and evaluate the robustness of discrimination in the presence of noise, we measured the effect of dataset dispersion on classification performance by training LDA classifiers to predict contact feature types from activation vectors in each speed group. (see Materials and Methods) and quantified classification rate. Overall, classification accuracy improved as contact speed decreased across noise conditions, with the consistently high classification accuracy (∼ >80%) observed at the slowest contact speed across the tested noise conditions (blue line in **Figure 4C**) Notably, this low speed closely matched the antenna movement speed of active touch during exploration (**Figure S3**). To maintain the same classification rate, noise would need to be limited to ∼8% and 3% at medium and high speeds, respectively. To ensure that our results were not specific to the classification method, we also trained a convolutional neural network (CNN) to classify contact features based on the tactile dataset. The results from using a CNN indicated that the classification rate improved as the contact speed decreased and thus consistent with those obtained using LDA classifier (**Figure S4**).

Taken together, for a fixed sampling rate, slower contact speeds could provide more contact time and spatial span with features, thereby providing more detailed contact information to facilitate feature discrimination. Collectively, our results support the hypothesis that contact speed is inversely proportional to tactile discriminability, with slower contact speeds, as observed in insects [17], enhancing the ability to distinguish tactile features by allowing for richer sensory integration, thus informing effective design strategies for robotic tactile sensing.

### Cockroach-inspired antenna stiffness gradients enhance tactile classification

Building upon the finding that contact speed affects tactile feature discrimination, we investigated how the mechanical properties of the antenna itself affect tactile field data dispersion and, consequently, tactile discriminability (**Figure 4E**). We hypothesized that the exponentially decreasing stiffness gradient observed in cockroach antennae enhances tactile discrimination by promoting a sparser and more dispersed tactile dataset. To test this hypothesis, we compared tactile field datasets generated using three distinct stiffness profiles: exponential (cockroach-inspired), linear decreasing, and constant (uniform stiffness) (**Figure S5**). To quantify data dispersion across these three models, we computed dispersion indices as done previously (Pearson correlation, SSIM, MSE, and NMI) within each group. The exponentially decreasing stiffness profile produced the lowest Pearson correlation, SSIM values, and NMI values, along with the highest MSE values, indicating greater dataset dispersion (**Figure 4B** and **Figure S2**). A Kruskal-Wallis test on NMI indicated a significant effect of stiffness profile on both activation dataset (*p* = 1.6e-2) and tactile field dataset (*p* = 2.5e-18). Post-hoc multiple comparisons showed that all pairwise differences between stiffness profiles on tactile field dataset were significant (*p*<0.05). The exponentially decreasing stiffness profile allows for more distributed bending along the antenna, generating a more diverse tactile dataset that enhances classification. We further analyzed the effect of stiffness profiles by applying cNMFsc encoding and evaluating entropy of tactile fields (see Methods). The bases derived from the linear decreasing and constant stiffness profile models exhibited considerably higher entropy (average entropies of 2.7 and 2.8, respectively), indicating less structured representation, compared to those from the exponential decreasing stiffness profile model (average entropy of 1.9; **Figure 4F,G**). This suggests that the exponential stiffness gradient can enhance encoding of tactile stimuli.

To demonstrate the impact of mechanical properties on the robustness of tactile classification, we trained LDA classifiers using slow-speed tactile field datasets generated under these three different stiffness profiles with increasing noise levels. The classification accuracy was highest for the exponential decreasing stiffness profile (black line in **Figure 4D**). Although the linear stiffness profile produced greater overall dataset dispersion than the constant stiffness profile (**Figure 4B**), their classification performances were less robust with accuracy decreasing as noise increased. This occurs because increased within-class variability in the linear stiffness profile outweighs gains in between-class separation, causing greater overlap in tactile feature space and reduced robustness. In contrast, the constant stiffness profile produces more compact tactile representations with lower within-class variance (**Figure S5**), making it more robust to noise than linear stiffness profile. Taken together, these results demonstrate that cockroach-inspired antenna mechanics substantially enhance tactile discrimination by generating more informative and structured tactile data compared to more conventional touch sensor stiffness profiles. This finding supports the biological hypothesis that mechanical properties such as stiffness gradients inherently simplify tactile information processing, offering insights for the development of physically intelligent robotic tactile sensors.

### Influence of mechanical parameters on tactile information encoding

To identify and generalize mechanical design principles governing tactile sensing performance, we systematically explored how antenna mechanical parameters influence tactile discrimination by mapping the NMI across a high-dimensional parameter space of tactile fields (**Figure 5**; see Materials and Methods). Lower NMI values correspond to greater dispersion of tactile fields across contact conditions, indicating improved separability of tactile features. We examined the effect of three parameters: exponential decay gradient (*p*_3_), normalized base stiffness 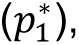 and damping-to-stiffness ratio (*c*_*n*_/*k*_*n*_), as indicators of mechanical factors that jointly governed tactile discrimination performance (**Figure 5A**). Across fixed damping-to-stiffness ratios, the lowest NMI (dark blue regions) emerged only within a specific combination of low base stiffness and steep exponential decay gradients. In general, reducing the base stiffness increased tactile signal dispersion by enhancing antenna compliance and deformation. However, this effect strongly depended on the spatial stiffness gradient. For example, antennae with relatively stiff bases maintained low NMI values when paired with sufficiently steep decay gradient (*p*_3_ = −0.1), suggesting that compliant distal regions can preserve distinct tactile fields despite increased proximal rigidity. Together, 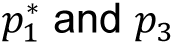 act as a dual-tuning mechanism: 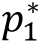 primarily regulates the global mechanical sensitivity of the antenna, whereas *p*_3_shapes the spatial distribution of that sensitivity along the antenna’s length.

**Figure 5.**
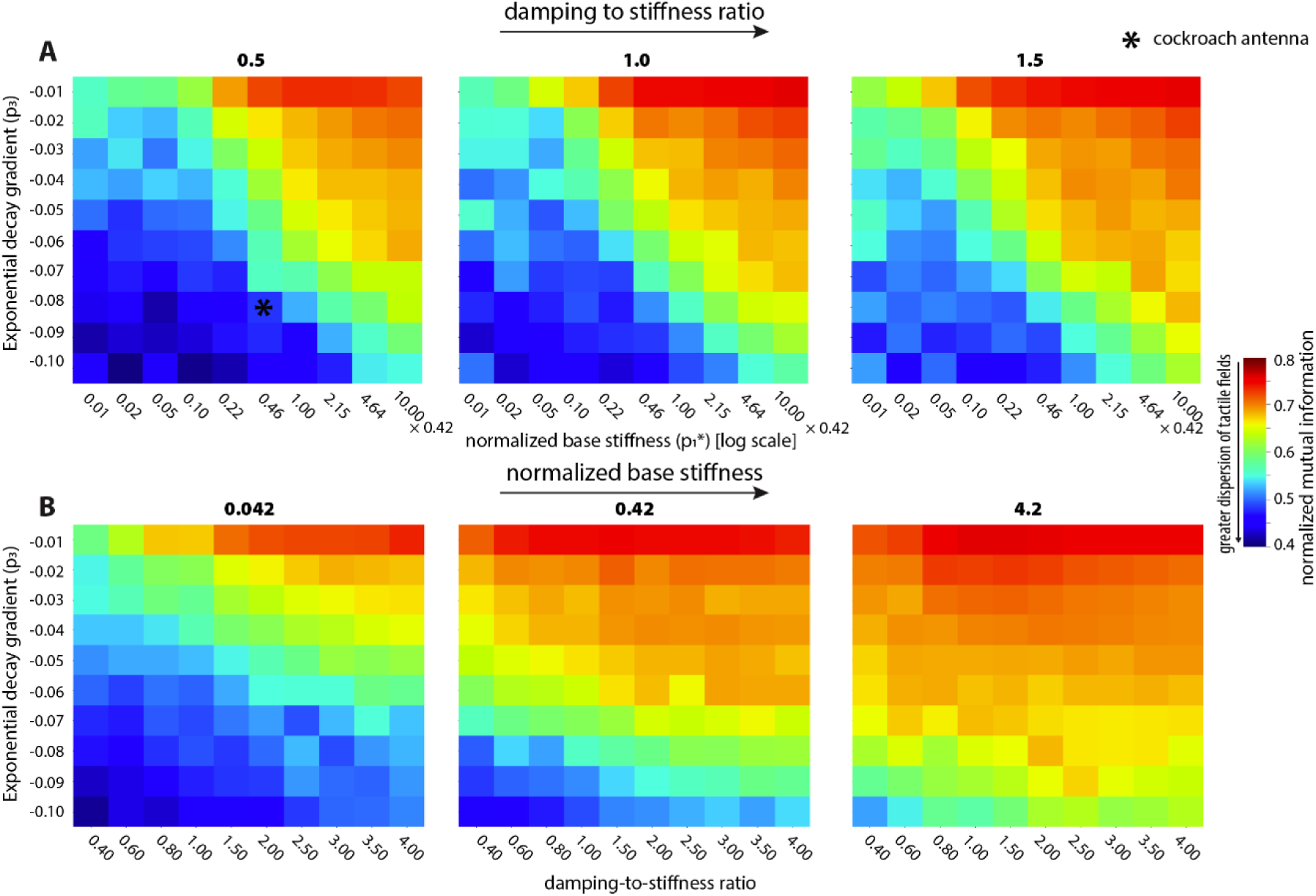
Mechanical parameter landscape governing tactile information encoding. (**A**) Heatmaps showing normalized mutual information (NMI) across parameter combinations of normalized base stiffness 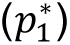 and exponential stiffness decay gradient (*p*_3_) for three damping-to-stiffness ratios (*c*_*n*_/*k*_*n*_= 0.5, 1.0, 1.5). *: approximate location of cockroach antenna 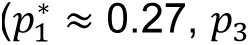 ≈ 0.27, *p*_3_ =-0.08, *c*_*n*_/*k*_*n*_ = 0.67 with the simplifying assumption that *p*_3_ = *p*_4_. **(B)** Heatmaps showing the influence of damping-to-stiffness ratio (*c*_*n*_/*k*_*n*_) and exponential stiffness decay gradient (*p*_3_) across three fixed normalized base stiffnesses 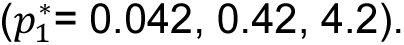

The damping-to-stiffness ratio (*c*_*n*_/*k*_*n*_) further structured tactile fields. The influence of damping was most pronounced in high-compliance regimes, like 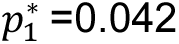 which is around 10 orders of magnitude smaller than 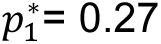 of the biological antenna model (**Figure 5B**). Lower damping-to-stiffness ratios produced lower NMI values but introduced high-frequency oscillations following the initial contact. In contrast, higher damping-to-stiffness ratios suppressed these oscillatory dynamics, generating smoother tactile fields that reduced dispersion across contact conditions. As a result, increasing damping generally increased NMI.

These results reveal a tradeoff between tactile discrimination and mechanics. Antennae with low base stiffness, steep stiffness gradients, and low damping-to-stiffness ratios (low 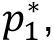 steep *p*_3_, low *c*_*n*_/*k*_*n*_) produced the most separable tactile fields, but such highly compliant antennae may also exhibit undesirable oscillations and reduced structural stability. Conversely, stiffer and more heavily damped antennae produced more mechanically stable responses at the cost of reduced tactile discriminability. Interestingly, the antenna of *P. americana* is approximately in a region with relatively low NMI and intermediate base stiffness and damping, suggesting that it is effective at tactile discrimination (**Figure 5A**). These findings suggest that “optimal” antenna design depends on balancing information-rich tactile encoding with the mechanical and operational constraints of the target robotic application.

### Simulation-generated tactile field principles transfer effectively to distributed soft robotic antenna designs

To demonstrate the practical applicability of our bioinspired tactile sensing framework and validate its generality and robustness against simulation-to-real gaps—such as contact model inaccuracies [30]—we transferred our computational findings to a physical robotic model. We hypothesized that the tactile encoding principles discovered through simulation—particularly the effectiveness of cockroach-inspired stiffness gradients and active touch behavioral strategies—could be successfully implemented in a physically intelligent, distributed robotic antenna. To demonstrate these concepts, we built a near-cockroach scale (compact: 7.3×1.6×0.2cm, lightweight: 491mg, low-power: 32mW) robotic antenna equipped with 8 capacitive hinge mechanosensors experiencing natural contact physics and capable of tactile feature discrimination (**Figure 6A**) [21]. Like the biological antenna, this flexible robotic antenna featured a linearly decreasing stiffness gradient and can resolve hinge angles (<1° resolution) at a maximum sampling rate of 1000 Hz, enabling the generation of high-fidelity tactile fields (**Figure 6A**). However, our robotic model is different (i.e. significantly simplified) to its biological analog in the following ways because of deliberate design choices. Due to manufacturing challenges, morphological variations such an exponential gradient or an increased number of mechanosensory units were not tested, but simulation of this eight-segment model yielded results consistent with the higher-degree-of-freedom model. However, this presents an interesting opportunity to leverage our simulation framework using the linear stiffness gradient with lower classification robustness (**Figure 4D**) and few mechanosensory units allowed us to demonstrate that even suboptimal design combinations inspired from the biological model could yield adequate tactile fields for performing effective feature classification in a majority of conditions. To confirm that the simulation environment effectively captured realistic antenna mechanics, we tuned the physics model with eight segments to closely match the robotic antenna’s impulse response (**Figure 6B**, **Movie S4**; see Materials and Methods). Taking advantage of the computational efficiency of the simulation environment, we generated a tactile field dataset encompassing two contact features, five contact locations, and three contact speeds. We then trained a support vector machine (SVM) classifier to distinguish these features (30 distinct tactile features for training, see Materials and Methods). To evaluate the sim-to-real transferability of our computational results, we applied the trained classifier directly to the robotic antenna in real-world contact experiments (**Figure 6C**). Real-world contact experiments were conducted by colliding the robotic antenna with ball and box features at its tip under three different speeds (6 distinct tactile features for testing). Using tactile fields generated from the sensory data of the robotic model and the SVM classifier trained on the simulation dataset, 83% of the experimental conditions were accurately predicted in terms of collision features, speed, and location (**Figure 6D,E**; **Figure S6**). In the misclassified class, the model correctly identified speed and location but failed to distinguish the feature, likely due to the limited dispersion in the dataset resulting either from the limited spatial resolution or simple (linear) stiffness gradient of the robotic antenna. Overall, these results validate the practical effectiveness and robustness of bioinspired tactile sensing principles—particularly stiffness gradients and active contact speeds—in realistic robotic implementations. Furthermore, this successful sim-to-real transfer underscores the feasibility of using our bioinspired tactile sensing framework as future work to create physically intelligent, distributed soft tactile sensors capable of enhanced perception in complex environments.

**Figure 6.**
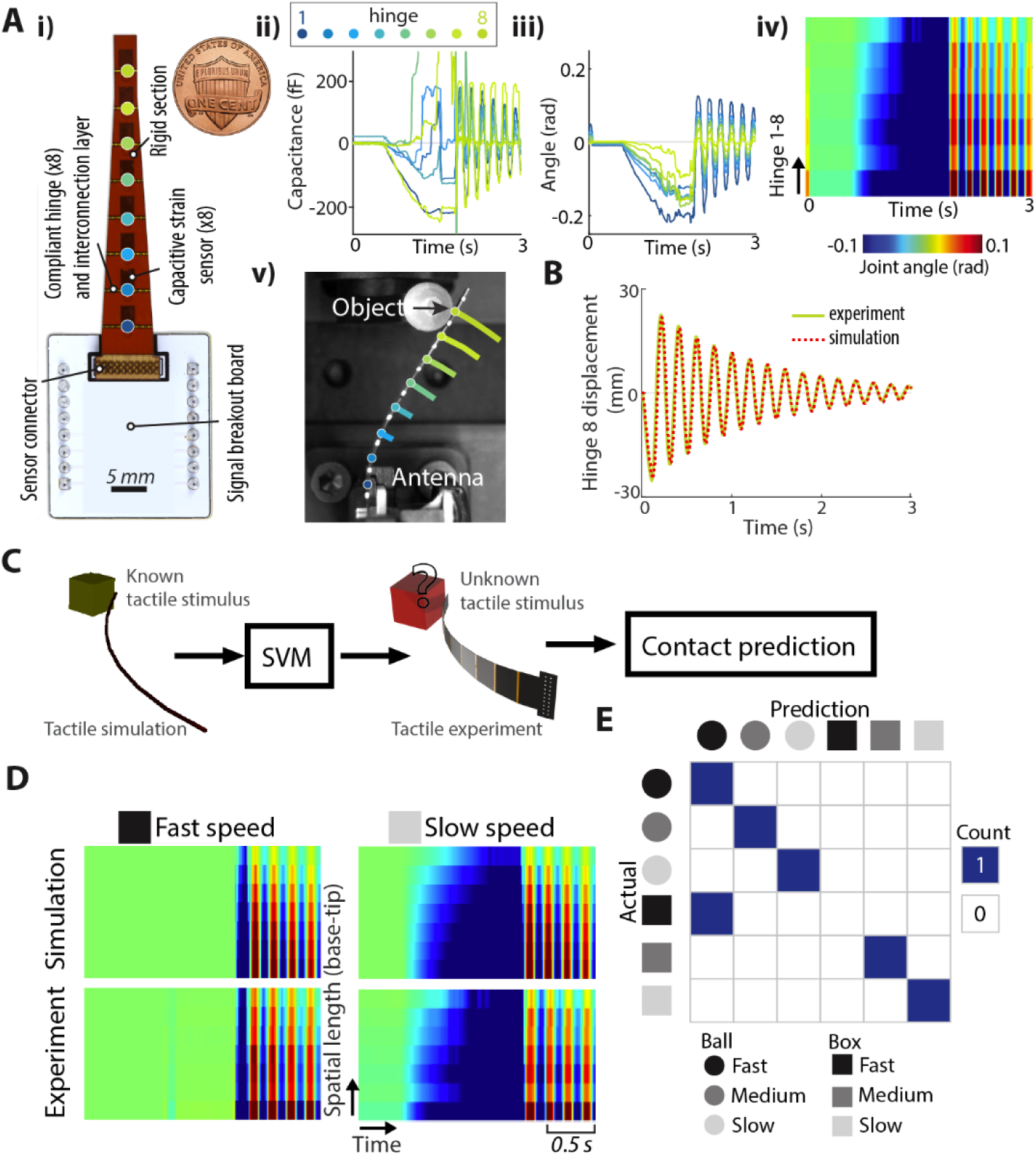
Transfer of classifier from simulation to robophysical antenna for tactile feature discrimination. (**A**) A distributed sensor system with eight embedded strain sensors. (**i**) Layout of the robotic antenna system; colored dots indicate the locations of sensors on the hinges. (**ii**) Raw capacitance readouts from the sensors during a collision scenario (collision at the tip with a speed of 0.027 m/s). (**iii**) Predicted hinge angles based on linear regression from sensor calibration [52]. (**iv**) Tactile fields corresponding to this collision scenario. (**v**) Ball collision at the tip with a speed of 0.027 m/s. (**B**) The impulse responses from the robophysical antenna and simulation model. (**C**) Transfer of a tactile field dataset with rich tactile stimuli from simulation to the distributed sensor system. (**D**) Two examples of tactile fields under the same condition: one from simulation and the other from sensory data collected by the robotic antenna. (**E**) Confusion matrix of the transferred SVM classifier. “Ball” and “box” indicate the two types of collision features. Black, gray, and white correspond to collision speeds of 0.270 m/s, 0.135 m/s, and 0.027 m/s, respectively.

## Discussion

Our study supports the hypothesis that physical intelligence—sensory processing embedded within mechanical structures—could represent an adaptive strategy in insect antenna [2,31]. By combining experiments, simulation, and physical modeling, our findings show that insect-inspired antennae mechanics, particularly non-linear stiffness gradients, together with active touch speed modulation, can simplify tactile feature discrimination by increasing data sparsity and dispersion by decreasing mutual information across tactile fields. Although demonstrated here through insect antennae-inspired sensors, such physical intelligence mechanisms could be broadly applied to other tactile modalities and robotic architectures, opening pathways for enhanced efficient tactile perception beyond simple navigation tasks (e.g., object handling, environmental exploration, and autonomous manipulation) in real-world complex environments.

### Advancing physically intelligent tactile perception inspired by insects

Our results pave the way toward a comprehensive design framework for physically intelligent tactile sensing inspired by insect antennae (**Figure 1**). First, analogous to cameras, distributed tactile sensor resolution is limited by the spacing between adjacent mechanosensors. The spacing between mechanosensors is inversely proportional to spatial resolution. Second, the sampling rate defines the temporal resolution to distinguish between different shapes, size, speed of contact, etc. Further, the integration time (sampling window; see below) influences tactile perception. Related to integration time is contact speed, as slower contact of the antenna with a feature will increase contact time and therefore the information available. Third, the flexural stiffness of the sensor must be sufficiently low to conform to tactile features. Specifically, a more compliant sensor increases the information available by activating more sensor elements, thereby sampling information from a larger region of space (**Figure 1A**). In addition to compliance, a stiffness gradient, as described in the cockroach antenna, can promote sparsity in tactile fields during feature contact, which can improve feature classification. Here, through detailed computational simulations and robophysical experiments, we provide quantitative support that physically intelligent mechanical designs significantly simplify tactile encoding, thus reducing computational and cognitive demands associated with tactile information processing.

Taken together, spatial resolution (sensor spacing), temporal resolution (sampling rate, integration time), and mechanics (compliance, gradient) must be considered to design effective tactile sensors (**Figure 1B**). Crucially, this work establishes the foundation for a broader tactile sensor design framework applicable to diverse robotic contexts beyond navigation alone. The demonstrated principles of mechanical stiffness gradients and active sensing have potential relevance across multiple robotic tasks, including manipulation of delicate objects, exploration of complex terrains, human-robot interaction, and prosthetic sensing systems. For example, robotic grippers employing similar bioinspired mechanical gradients could achieve improved sensitivity, dexterity, and adaptability, enabling reliable manipulation of diverse objects in uncertain environments [32,33]. Similarly, wearable tactile sensors incorporating these principles could significantly enhance feedback quality and intuitive control in prosthetic and assistive technologies.

### Trade-off between speed of contact (energy) and information

Temporal integration is a fundamental feature in perceptual decision-making [12], allowing sensory systems to accumulate information over time. Longer sensing durations enable greater temporal integration, improving sensory estimates. The speed-accuracy tradeoff (SAT) is a well-established principle in psychophysics, describing how animals and humans balance rapid decision-making with the need for precise sensory processing [13,34]. In visual perception, acuity varies across the visual field, and humans acquire high-resolution information through brief fixation periods separated by saccades. Studies on rapid serial visual presentation have shown that detection accuracy improves with longer exposure times, highlighting the importance of temporal integration in visual discrimination [35]. Similarly, in tactile perception, temporal cues play a crucial role in spatial-frequency discrimination, such as detecting surface textures [36]. These consistent observations suggest a generalized biological principle of active sensory modulation, offering valuable insights for adaptive robotic sensing strategies. Moreover, this SAT principle aligns with our findings, where slower contact speeds resulted in greater dataset dispersion, improving the classification of tactile features. Taken together, our findings provide a link between contact speed and tactile dataset dispersion in both simulated and experimental settings. Distributed soft sensor systems, such as the robotic sensor developed here, could optimize tactile information processing by dynamically adjusting contact speed. Future work could explore adaptive speed control strategies in robotic systems, allowing for dynamic modulation of contact speed based on task demands, similar to active touch sensing behaviors in animals.

### Role of mechanics in tactile discrimination

Mechanics and sensing are tightly coupled in biological systems [31]. For instance, insect wings—which act like actuators and sensors—can enhance body rotation detection with their nonuniform stiffness gradient [19]. The tapering of whiskers enhances the reliability of tactile information by facilitating stick-slip motions during surface exploration, which are essential for texture discrimination. In contrast, cylindrical whiskers tend to become stuck or immobilized when sweeping across fine textures, which reduces sensory effectiveness [20]. While these sensors have demonstrated strong performance in spatial mapping and texture discrimination tasks [37,38], their capabilities are often limited by structural rigidity and limited sensor density. In contrast, insect antenna-like sensors offer inherent advantages through their mechanical gradients and high-density sensor distribution. Our findings related to exponential stiffness gradients in cockroach antennae raise the intriguing possibility that these mechanical properties represent convergent evolutionary solutions across tactile sensory systems in diverse taxa. Indeed, mechanical gradients and sensor distributions similar to those we observed have been documented in diverse arthropods including stick insects, crickets and locusts [39,40]. Likewise, crayfish antennae exhibit stiffness gradients enabling precise water-flow detection and olfaction [41]. Additionally, the homology between antennae and legs extends to the underlying sensorimotor system [42] hinting at potential mechanical gradient-based advantages during force encoding and control [43] during other tactile behaviors such as legged locomotion. Future comparative studies quantifying stiffness gradients, sensor densities, and segmental dimensions across diverse species could provide compelling quantitative support for tactile sensing convergence, offering deeper insights into evolutionary biomechanics and sensing strategies.

### Role of stiffness gradients and damping in tactile discrimination

To begin to understand how mechanics shape tactile fields, we treated the antenna as a multi-link system with the cockroach-inspired mechanics, governed by normalized base stiffness 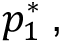 exponential gradient *p*_3_, and damping-to-stiffness ratio (*c*_*n*_/*k*_*n*_). We mapped the NMI across this parameter space and confirmed that tactile fields are influenced jointly by these parameters. While 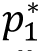 determines the global mechanical sensitivity of the structure, *p*_3_modulates the spatial distribution of deformation along the antenna length. A more compliant base paired with a steep gradient generally allows the antenna to produce a greater dispersion of tactile fields, which in turn improves tactile discrimination. However, this advantage comes at the cost of structural stability. Consequently, our results suggest that the’optimal’ antenna design is application dependent: high-speed robotic navigation may favor stiffer, more heavily damped configurations, whereas complex object classification tasks benefit from increased tactile field dispersion provided by softer, less damped, and highly tapered mechanical profiles.

### Implications for understanding the sense of touch in biology

The tactile dataset generated in this study illustrates the dynamic variations in tactile stimuli encountered during antennal contact. At present, it remains unknown whether cockroaches can actively discriminate between these stimuli. To explore this possibility, we applied cNMFsc and encoding of tactile datasets, to mimic neural signal processing in the antenna. This encoding provides a possible characterization of how tactile signals may be processed within the cockroach’s sensory systems. Notably, the different bases extracted through cNMFsc preserved information about the contact region along the antenna, suggesting the spatial organization of these bases may reflect a form of topographic mapping of tactile feature. This topographic mapping is commonly observed in biological systems, such as somatotopic maps in the somatosensory cortex [44] and visual feature maps [45]. In cockroaches, adjacent sensory receptors along the antenna could project to adjacent regions in the brain, preserving spatial relationships [46,47]. Our spatiotemporal encoding model provides support for this structure. Specifically, the temporal activation patterns reflect a progression from proximal to distal regions of the antenna, mapping contact location into basis space. This topographic-like representation may facilitate efficient neural decoding of contact location, supporting efficient tactile localization.

### Opportunities for bio-inspired robotics

Our work provides a framework for designing compliant robotic tactile sensors. While whisker-inspired tactile sensors have been widely explored [3], distributed soft sensors inspired by insect antennae offer unique advantages not afforded by whisker sensors [48]. In contrast to whiskers, many insect antennae are multi-segmented, continuously deformable structures with distributed sensing along their length [49,50]. This structural complexity could allow for rich information extraction and broad contact-based perception from the environment. Optimizing the structural and mechanical properties of bio-inspired antennae can provide novel opportunities for robotic perception and interaction with complex environments [6,48].

These design principles also translate to physical systems. One key advantage of our bio-inspired antenna system is its ability to discriminate environmental features while maintaining low energy consumption and minimal weight. While many conventional robotic tactile sensors—such as piezoelectric or resistive tactile sensors—can require supporting electronics with relatively high-power demands, our bio-inspired distributed soft sensors are based on capacitive sensors, which is well known for its low energy requirements [51]. Our robotic antenna operates with a total power consumption below 60 mW and weighs under 500 mg, making it ideal for energy-efficient and lightweight applications [52]. This level of efficiency is particularly beneficial for autonomous insect-scale robotic systems operating in resource-constrained environments.

## Materials and Methods

### Bio-inspired antenna model to simulate the transmission of tactile stimuli

Inspired by the morphology and mechanics of the antenna (flagellum) of *P. americana* [15,28], our model included a sequence of 140 rigid links connected by hinge joints (**Figure 1D**). The radius of each link was kept constant, and the mechanical properties of each joint were modeled by modulating the link mass, hinge stiffness and hinge damping. The length of each segment increases from the base to the tip in the antenna of the American cockroach [15,53]. This variation was modeled using a sigmoid function fitted from experimental data (squared norm of the residual = 8.7e-5, see **Figure S7**)[15].

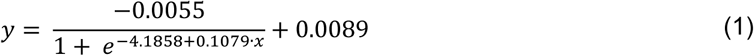

where *y* represents the proportion of the length of the segment to the total length, and *x* is the ratio of the segment number to the total number of segments.

The mass of each segment was described by the following equation, where the assumption was that the density is constant along the antenna, resulting in the mass following the same exponential decreasing relationship fitting of the square of the radius [15].

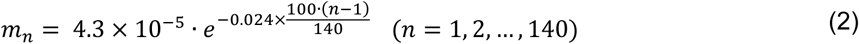

where *m*_*n*_ (g) is the mass of the *n*^th^ segment.

The stiffness *k* and damping *c* of each joint *n* were described by exponential functions modeling the mechanics from base to the tip of the antenna [15].

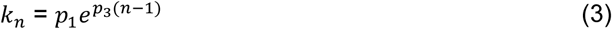

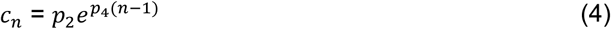

where *p*_1_ and *p*_2_ are the stiffness and damping constants at the first (base) joint, respectively. The rates of exponentially decrease in stiffness and damping of joints along the length are represented by *p*_3_and *p*_4_, respectively. To compare the influence of different stiffness and damping profiles, a linear decreasing and constant stiffness and corresponding damping profiles were also implemented in the antenna model.

Stiffness and damping parameters for each function were optimized using nonlinear methods to match the damped natural frequency and damping ratio obtained from step deflection data (**Figure 2B**) [15]. Additionally, the deflection ratios between the middle regions of the antenna (approximately 71% and 86% of its full length) and the tip (100% of full length), obtained from impulse collision experiments (**Figure 2B**), were used as target quasistatic constraints in the optimization [15]. We used normalized Euclidean distance to quantify the total error of the simulations across the two scenarios, as calculated using

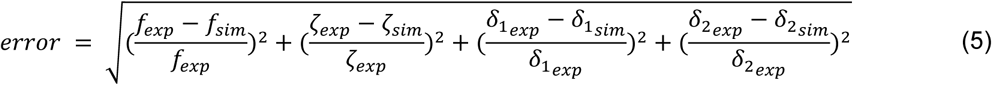

with constants defined in Table 1.

**Table 1.** Dynamic and static properties from experiments [15].

| Damped natural frequency<br>(Hz) | Damping<br>ratio | The deflection ratio between the middle point<br>and tip at the maximum deflection time. |  |
| --- | --- | --- | --- |
| $f_{exp} = 18$ | $\zeta_{exp} = 0.52$ | $\delta_{1exp} = 16.5 \%$ | $\delta_{2exp} = 45.1 \%$ |

To handle non-differentiable target functions, a nonlinear Powell’s method was used to minimize the error [54]. After optimization, the parameter values were determined as follows: *p*_1_ = 37.78, *p*_2_ = 25.24, *p*_3_ = −0.0827, and *p*_4_ = −0.0533, resulting in a local minimum error of 0.067 (**Movie S1**). It is worth noting that different combinations of *p*_1_to *p*_4_ can lead to similar errors. As a comparison, the median symmetric accuracy (MSA), as previously used [55], was 0.0035. The MSA is more robust to outliers as it relies on the median of the target values, whereas the normalized Euclidean distance gives equal weight to each target.

### Generation of tactile fields from contact

We implemented the antenna model using the MuJuCo physics engine [56], chosen for its efficiency and accuracy in simulating multi-rigid body dynamics in contact-rich environments. The default numerical integrator, the implicit-in-velocity Euler method, was employed with pyramidal contact cone in MuJoCo’s soft contact model. This model relaxes the strict complementarity constraint in linear complementarity formulation to increase physical realism for soft contacts [57].

We recorded joint angles as the antenna contacted distinct features at distinct locations and contact speeds. We defined a “tactile field” as the full set of segment curvature (or joint angle) across the antenna in time during a tactile event. Thus, each tactile field *a*(*x*, *t*) has dimension *x*, the distance to the base of the antenna, and time *t* (**Figure 2E**). To generate a comprehensive tactile dataset encompassing a set of mechanical stimuli applied to the antenna, we simulated three different contact speeds: 0.6 m/s (cockroach evasive speed [29]), 0.3 m/s (medium speed), and 0.06 m/s (antenna slow exploration speed [58], see Supplementary Methods). These speeds were chosen to represent a range of antenna-mediated behaviors. We tested collisions with six different features at five locations along the antenna (indicated by red dots in **Figure 2D**), spanning the middle to the tip region to reflect a broad set of contact points. Joint angles from 56 points, representing the locations of campaniform sensilla (strain sensors) along the antenna length [46], were recorded every 1 ms, resulting in a dataset of 90 tactile fields (**Figure S5**).

### Spatiotemporal encoding of tactile fields

To extract features from tactile fields by dimensionality reduction, we performed spatiotemporal encoding on the tactile fields based on convolutive non-negative matrix factorization with sparsity constraints (cNMFsc), which has been previously used to encode tactile stimuli to the human hand [59]. cNMFsc decomposed the tactile stimuli, *a*(*x*, *t*), into a set of spatiotemporal basis sequences *w*_*i*_(*x*, *t*), and their corresponding activation vectors ℎ_*i*_(*t*). The algorithm minimizes the L2 norm between *a*(*x*, *t*) and the estimated tactile stimuli,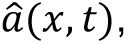 subject to the sparsity constraints [60] on the activation vectors:

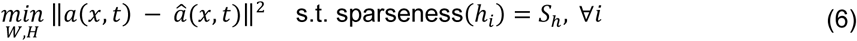

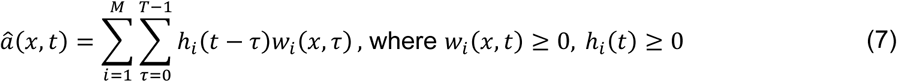

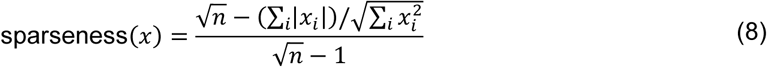

where 0 ≤ *S*_ℎ_ ≤ 1 is the defined sparseness, *n* is the dimensionality of vector *x* and *M* is number of bases. The sparseness was chosen as 0.8 in this study. We set the duration of the spatiotemporal basis, *T*, to 40 ms, which approximates the cockroach’s response time to impulse-like perturbations of the antenna [16,29,61], although the spatiotemporal profile of the bases exhibited little variation across different durations *T* (**Figure S8**).

### Dataset dispersion evaluation

To assess the similarity between tactile fields, we computed the Pearson correlation coefficient, structural similarity index (SSIM), mean squared error (MSE), and normalized mutual information (NMI) for each tactile field against all other tactile fields in the dataset. For each tactile field, we identified its closest match based on these metrics, selecting the maximum value of Pearson correlation coefficient, SSIM, NMI, and the minimum value of MSE. This process was repeated for all tactile fields to quantify the most similar counterpart for each field in the dataset. Finally, the average Pearson correlation, SSIM, MSE, and NMI across all comparisons were used to evaluate the overall dispersion of the dataset.

### Entropy and normalized mutual information analysis

To quantify the information content and statistical dependence of tactile signals, we computed Shannon entropy and NMI using histogram-based estimators. For a given scalar signal *X* (e.g., elements of the tactile field or activation coefficients), the Shannon entropy *H*(*X*) was estimated as:

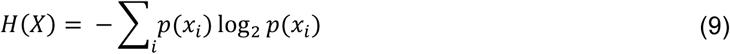

where *p*(*x*_*i*_) represents the probability of observations falling within the *i*-th bin of a histogram-based discretization. Signals were discretized into 20 bins, and probabilities were estimated using normalized histograms.

To quantify the statistical dependence between two signals *X* and *Y*, we computed their normalized mutual information:

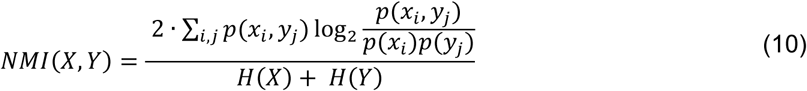

where *p*(*x*_*i*_, *y*_*j*_) is the joint probability estimated using a two-dimensional histogram, and *p*(*x*_*i*_), *p*(*y*_*j*_) are the marginal distributions.

### Discrete multilink model with stiffness scaling and parameter space exploration

To further study the influence of exponential decreasing gradient in tactile field dataset dispersion to infer tactile discrimination, we normalize our multilink model to explore how the parameters *p*_1_ and *p*_3_ in Equation (3) change the NMI of tactile fields across simulation. The system can be modeled as a discrete multilink system consisting of *N* rigid segments connected by *N* rotational hinges. To ensure that the mechanical properties of the simulated antenna remain consistent regardless of the discretization resolution or total physical size, the hinge stiffness *k*_*n*_ is derived from the flexural stiffness *EI* of a continuous beam [62]. The stiffness of an individual hinge is defined as:

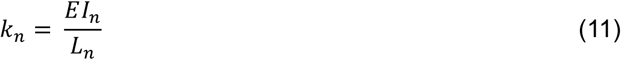

where *L*_*n*_ represents the length of the *n*^th^ segment. For an antenna of total length *L*, it is divided into *N* segments of uniform length 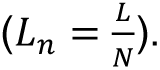 According to Equation (3), we have:

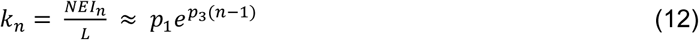

where 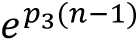 describe the exponentially decreasing second moment of area *I*_*n*_. Thus, from the discretization perspective, the base stiffness parameter *p*_1_ scales proportionally with the number of segments:

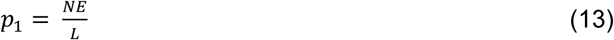

From the structural perspective, bending is driven by the moment *M* and the shape is defined by curvature *κ* and according to Equation (13):

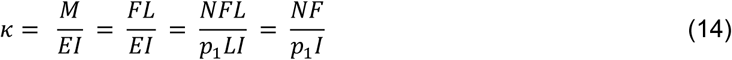

Thus, it is also found that the base stiffness parameter *p*_1_ scales proportionally with the number of segments *N* regardless of the antenna length *L* to ensure that the model preserves its characteristic bending shape across different scales. Overall, we can normalize the base stiffness parameter *p*_1_ by

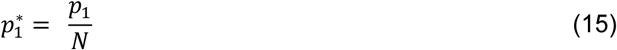

In order to generalize the parameter exploration space, we choose the model length *L* to be 42 mm which is the same as the biological antenna model used previously and discretized it into 100 segments. To reduce dimensions, we define a damping-to-stiffness ratio (*c*_*n*_/*k*_*n*_). To evaluate the antenna mechanical response across parameters, a dataset was generated consisting of 30 distinct collision conditions using the same method as described previously. For each condition, the time-series data of 100 hinge sensors were recorded over a simulation duration of 1000 ms. The NMI evaluation metric was used as described previously.

Two systematic grid searches were conducted to identify the optimal mechanical configuration for tactile discrimination:

1. Stiffness-gradient influence: we explored the base stiffness 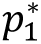 across a logarithmic scale from 0.0042 to 4.2 and the exponential decay *p*_3_ from-0.1 to-0.01, given that the 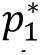 and *p*_3_ of the optimized biological antenna is 0.27 and-0.0827. This sweep was repeated for 3 damping ratios (*c*_*n*_/*k*_*n*_ ∈ {0.5, 1, 2}).
2. Damping-Gradient Influence: We investigated the damping-to-stiffness ratio *c*_*n*_/*k*_*n*_ across a range from 0.4 to 4.0 and the exponential decay *p*_3_ from-0.1 to-0.01. The range of damping ratio is determined by the optimized biological antenna. This exploration was performed across three base stiffness levels 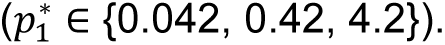.

### Feature classification from tactile fields

To further quantify the influence of tactile field dispersion on tactile discrimination, we designed a classification task with the objective of using the activation vectors to identify contact locations and features (**Figure S9**). First, we performed Principal Component Analysis (PCA) on the flattened activation vector and selected the first 30 features, which could capture ∼90% of the variance for the test sets. To simulate real-world noise and augment the data, we added Gaussian noise (*μ* = 0) to the activation vectors and applied dimensionality reduction through the same PCA process. The noise ratio was defined as the deviation of Gaussian noise relative to the maximum value of each activation vector. We initially avoided complex classification methods that would require extensive model tuning, such as convolutional neural networks (CNNs), although classification results using CNNs yielded similar conclusions (**Figure S4**). Instead, we chose Linear Discriminant Analysis (LDA) classifiers, which provides a parsimonious method. The data augmentation, dimensionality reduction, and training steps were iterated ten times to derive the average accuracy of the test sets. Collectively, the average accuracy of the test sets provided a measure of tactile discriminability.

## Statistics

We report mean ± 1 standard deviation, unless otherwise specified. Statistical significance was determined via Kruskal-Wallis with post-hoc test; ***: p < 0.001, **: p < 0.01, *: p < 0.05, n.s.: p > 0.05.

## Data accessibility statement

All data and code are available on Penn State ScholarSphere: https://doi.org/10.26207/tmsg-ad78

## Acknowledgments

We thank Netta Cohen for helpful discussions. Research was sponsored by the Army Research Office and was accomplished under Grant W911NF- 23-1-0039 (JMM, KJ) and by the Alfred P. Sloan Research Fellowship FG-2021-16388 (JMM).

